# Diversity and ecological function of urease-producing bacteria in the cultivation environment of *Gracilariopsis lemaneiformis*

**DOI:** 10.1101/2022.12.15.520688

**Authors:** Pengbing Pei, Hong Du, Muhammad Aslam, Hui Wang, Peilin Ye, Tangcheng Li, Honghao Liang, Zezhi Zhang, Xiao Ke, Qi Lin, Weizhou Chen

## Abstract

Urease-producing bacteria (UPB) provide inorganic nitrogen for primary producers by hydrolysis of urea. They play an important role in marine nitrogen cycle. However, there is still incomplete understanding of UPB and their ecological functions in the cultivation environment of red macroalage *Gracilariopsis lemaneiformis*. This study comprehensively analyzed the diversity of culturable UPB and explored their effects on urea uptake by *G. lemaneiformis*. Total 34 isolates belonging to four main bacterial phyla i.e. Proteobacteria, Bacteroidetes, Firmicutes, and Actinobacteria were identified through 16S rRNA sequencing and were screened for UPB by urea agar chromogenic medium assay and *ureC* gene cloning. Our data revealed that only 8 strains were found containing urease. These all UPB exhibited different urease activities by Berthelot reaction colorimetry assay. Furthermore, UPB with highest urease activity was selected to use as co-culture with *G. lemaneiformis* to explore its role in terms of promotion or inhibition of nitrogen uptake by *G. lemaneiformis*. The results showed that the urea consumption in culture media and the total cellular nitrogen in *G. lemaneiformis* found increased significantly in the UPB-co culture group than control i.e. in the sterile group (*p* < 0.05). Similarity, isotopic assay revealed that δ^15^N contents of *G. lemaneiformis* was significant higher in the UPB-co culture than in the control where δ^15^N-urea was the only nitrogen source in the culture media, indicating the UPB helped *G. lemaneiformis* to absorb more nitrogen from urea. Moreover, the highest content of δ^15^N was found in *G. lemaneiformis* with epiphytic bacteria, as compared to sterilized (control) showing that epiphytic bacteria along with UPB have compound effects in helping *G. lemaneiformis* absorb more nitrogen in urea. Taken together, these results provide unique insight into the ecological role of UPB and suggest that urease from macroalgae environment-associated bacteria might be important player in the marine nitrogen cycling.

**Importance:** To the best of our knowledge, this is the first study ever conducted to isolate the culturable UPB from the cultivation environment of *G. lemaneiformis* by urea agar chromogenic medium assay, and also evaluate the effects of UPB on urea utilization in *G. lemaneiformis* by stable isotopic tracer technique. This study provides a new insight into the mechanism of organic nitrogen uptake and utilization in *G. lemaneiformis*, and is of great significance for a better understanding of the ecological role of functional bacteria (e.g. urease-producing bacteria) in the marine nitrogen cycling.

## 1. Introduction

Nitrogen (N) is not only essential element of all life forms, but also one of the pivotal macronutrients that restrict biomass and primary productivity in the marine ecosystem (1, 2). N compounds in marine environment mainly exist in inorganic and organic forms (2, 3). In marine environment, the algae and marine plants use inorganic N directly (4, 5). However, organic N sources can also be utilized by the algae and marine plants when the amount of inorganic N in seawater is lower or even scarcer (6). Organic N mainly includes proteins, amino acid/sugars, nucleic acids, urea, and humic substances (3). Urea being smaller nitrogenous compound (7), is reported higher in concentration in the coastal ecosystems (8). It is has been observed that marine plants (e.g. seagrass) (9, 10) and algae (e.g. phytoplankton and macroalgae) (11, 12) use urea for their N source. However, for the absorption of organic N, both marine plants and algae need the assistance of microorganisms (13–15). It is found that microbiota associated with seagrass leaves increase N uptake availability of seagrass by mineralizing amino acids (14). Moreover, epiphytic bacteria of algae have found to promote algal growth by providing N nutrients (16). The source of N nutrients mainly comes from the N_2_ fixation and the decomposition of organic matter, both of which are performed by specific functional microorganisms (17, 18).

UPB are a group of functional microorganisms containing urease gene (*ureC*), which are involved in the conversion process of organic nitrogen into inorganic nitrogen, and therefore, play critical role in nitrogen cycling of marine environment (19, 20). It is urease produced by UPB that plays the function of organic nitrogen decomposition, which is widely existed in various bacterial genera, such as *Helicobacter, Proteus, Enterobacter, Pseudorhodobacter, Streptococcus, Escherichia, Lactobacillus, Enteroccocus, Weissella, Mycobacterium* (21–23). At present, there are some studies on urease in marine environment, both from the phylogenetic level to understand the diversity of UPB, and from the genetic level to explain the role of urease in urea utilization. For example, Siegl et al. (24) identified a 10 ORF containing urease gene cluster from the *Poribacteria* using single-cell genomics sequencing, including various ABC-transporter, three urease subunits *ureA, ureB, ureC*, as well as three accessory proteins. Another study provided the first insight into the bacterial potential in urea utilization by detecting the transcriptional activity of *ureC* gene as well as the phylogenetic diversity of bacteria with *ureC* gene (19). Although studies on urease have been carried out extensively in marine environments. However studies on UPB and the ecological functions of urease in cultivation environment of macroalgae are still unknown.

*G. lemaneiformis*, a member of the genus *Gracilariopsis*, is an important economic macroalgae and widely distributes in coastal areas of China (6, 25). It is currently the third largest mass cultivation seaweed in China, only after *Saccharina* and *Pyropia* (6), with a cultivation area reaching ca. 10,459 ha in 2020 (data from China Fishery Statistical Yearbook). The large-scale commercial cultivation of *G. lemaneiformis* has gradually formed an influential seaweed field, which has produced a series of ecological effects (6). For instance, it not only effectively improves the water quality of coastal environment, but also contributes to increase the marine carbon sink and mitigate climate change (26–28). Significant number of studies have been conducted with mainly focusing on the diversity of epiphytic bacterial community on *G. lemaneiformis*, the microbial community of seawater and sediment in the cultivation environment of *G. lemaneiformis* (18, 29). However, the diversity of functional bacteria in the cultivation environment of *G. lemaneiformis* have not been thoroughly studied. Our previous study reported five genus of UPB (such as *Lactobacillus, Escherichia-Shigella, Mesorhizobium, Helicobacter*, and *Streptococcus*) in the epiphytic bacterial community of *G. lemaneiformis* (18). It is found that, due to the high productivity and the ability to absorb large quantity of N (25), *G. lemaneiformis* confronts with the stress of inorganic N deficiency at the end of cultivation periods. Therefore, organic N (e.g. urea) as a supplementary nitrogen source. Moreover, *G. lemaneiformis* is found to utilize urea as a nitrogen source to maintain its growth (7), but the role of functional bacteria found in the cultivation environment of *G. lemaneiformis* are poorly understood. Therefore, our goal was to explore the diversity and urease activity of UPB in the cultivation environment of *G. lemaneiformis* along with their role in helping urea uptake by macroalgae and marine N cycle which is, to the best of our knowledge have not been attempted previously.

## 2. Materials and methods

### 2.1. Study area and sample information

The commercial seaweed cultivation area of Shen’ao Bay, Nan’ao Island is large-scale cultivation of *G. lemaneiformis* which is often accompanied by small scale cultivation of *Porphyria haitanensis*. The samples of *G. lemaneiformis, P. haitanensis*, and seawater were collected from algae culturing field of Nan’ao Island (117°6′40″E, 23°29′9″N), Shantou, Guangdong province, China. The algal samples were stored into sterile polyethylene bags containing surrounding seawater and the seawater samples were stored into sterile polyethylene bottles. Both algal and seawater samples were stored at 4 _o_C and transported to the laboratory for isolation of epiphytic and free-living bacteria.

### 2.2. Isolation of epiphytic and free-living bacteria

In the laboratory, *G. lemaneiformis* and *P. haitanensis* samples were washed three times with autoclaved seawater to eliminate loosely attached epiphytes, sand particles and other attached settlements (30, 31). After rinsing, firmly attached epiphytic bacteria were swabbed with sterile cotton buds and then were swabbed on marine agar 2216 E plates. For free-living bacteria, a 0.1 mL seawater sample was dropped onto marine agar 2216 E plates and spread with spreader. Plates were incubated at 28℃for 3 days. Morphological i.e. (size, shape, color etc) different bacterial colonies were picked with inoculating loop to make streak plates for getting purified colonies. This step was repeated twice in order to obtain pure individual colony. These pure colonies obtained were preserved at -80℃in marine broth supplemented with 25% sterile glycerol.

### 2.3. DNA extraction and 16S rDNA gene sequencing

The DNA of pure bacterial colonies was extracted by following the procedures described in the DNA extraction kit (TIANGEN Biotech, Beijing). The 16S rRNA gene was amplified using universal primers 27F (5’-AGAGTTTGATCMTGGCTCAG-3’) and 1492R (5’-TACGGYTACCTTGTTACGACTT-3’). PCR was carried out in 30 μL reaction mixture containing 1 μL genomic DNA, 3 μL Buffer, 3 μL of each primer, 2 μL dNTP, 0.2 μL DNA Polymerase (Sigma), and 17.8 μL H_2_O under thermal cycle of 95℃for five min, 35 cycles of 30 s at 95℃, 30 s at 55℃and 1 min at 72℃, followed by 72℃for 10 min in a Bio-rad T100_TM_ Thermal Cycler (Bio-Rad, USA). The quality of PCR products was verified by 1% agarose electrophoresis gel. The 3730XL DNA Analyzer (ABI, USA) was used for sequencing of pure isolates. The sequence data reported in this study have been deposited in the NCBI GenBank database under the accession number SUB11205669. All data are available at: https://submit.ncbi.nlm.nih.gov/subs/?search=SUB11205669.

### 2.4. Identification of UPB and urease gene (*ureC*)

The UPB were screened using urea agar chromogenic medium (10 mL), which contains 0.1 g tryptone, 0.1 g D-(+)-glucose, 0.2 g KH_2_PO_4_, 0.0012 g phenol red, 1.5 g agar, 2% urea solution (w/v), 0.1 L artificial seawater (ASW) (19). A 0.1 L ASW used in this study includes 0.11 g CaCl_2_, 1.02 g MgCl_2_·6H_2_O, 3.16 g NaCl, 0.075 g KCl, 0.1 g Na_2_SO_4_, 0.24 g Tris-HCl, 0.002 g NaHCO_3_, 0.1 L dH_2_O (pH = 5.87). The fresh isolates were streaked on urea agar chromogenic medium and cultured into incubator at 28 ℃for two days. Then the isolates were defined as UPB if the medium color changed from pale orange to pink or fuchsia.

The *ureC* gene of UPB was amplified using universal primers L_2_F (ATHGGYAARGCNGGNAAYCC) and L_2_R (GTBSHNCCCCARTCYTCRTG) (19). PCR was carried out in 20 μL under thermal cycle of 98℃for 30 s, 35 cycles of 10 s at 98℃, 30 s at 59℃and 30 s at 72℃, followed by 72℃for 2 min. PCR products were verified by 2% agarose electrophoresis gel and subjected to sequencing by using a 3730XL DNA Analyzer (ABI, USA).

### 2.5. Urease activity of UPB

A 2 mL UPB bacterial cultures inoculated in 100 mL fermentation medium at 28℃were cultured at 180 rpm for 24 h. The fermentation medium used in this study contains 0.2 g D-(+)-glucose, 0.1 g tryptone, 0.05 g yeast extract, 0.05 g beef extract, 0.02 g KH_2_PO_4_, 0.05 g NaCl, 0.5% urea, 0.005% Ni(NO_3_)_2_, 0.1 L ASW. Then the fermentation medium was centrifuged at 10,000 *g* for 20 min to separate bacteria. The bacterial precipitates were suspended in 5 mL PBS buffer, and mechanically homogenized using ultrasonic cell pulverizer (30 W, 4 s/4 s) for 3 min on ice. Next, the supernatant was collected at 12,000 *g* for 5 min at 4℃. The protein concentration of the supernatant was determined by Coomassie brilliant blue kit (A045-2-2, Nanjing Jiancheng Bioengineering Institute, Nanjing, China).

The urease activity was measured by Berthelot reaction colorimetry (32). Briefly, a 2.5 mL 10% sterile urea solution, a 5 mL PBS buffer (8.00 g NaCl, 0.20 g KCl, 1.44 g Na_2_HPO_4_, 0.24 g KH_2_PO_4_, pH=7.4) and 10 uL crude enzyme solution were mixed thoroughly and incubated at 40℃for 20 min. Then, 50 μL reaction solution, 400 μL sodium phenol solution, 300 μL 0.9% sodium hypochlorite solution were sequentially added to the reaction tube and incubated at 40℃for 20 min. Finally, the OD_578_ value of color-stabilized reaction solution was determined within 1 h. The crude enzyme solution was replaced by PBS buffer of equal volume, and the control experiment was carried out according to the same operations. Definition of enzyme activity U: 1 U was equal to the release of 1 μmol NH_3_ after 1 mg enzyme solution reacted at 40℃for 1 min.

### 2.6. Co-culture of *G. lemaneiformis* and UPB

*G. lemaneiformis* was washed with sterile seawater and pre-cultured in modified f/2 media for one week according to our previous study (33). After pre-culture, *G. lemaneiformis* was cultured in modified f/2 media with low nitrogen concentration (5.49 μmol·L^-1^) for four days. Prior to the experiment, the nine groups of *G. lemaneiformis* weighting 5.0±0.05 g each was subjected to strict bacterial removing following previous report with minor modifications (34, 35). Briefly, *G. lemaneiformis* was immersed in 2 L modified f/2 medium for 4 h and sonicated in a SCIENTZ JY92-ⅡN Sonicator (Ningbo Scientz Biotechnology Co., Ltd. Ningbo, China) for 10 min (50W, 5 s/5 s). Then, *G. lemaneiformis* was incubated in 2 L modified f/2 media containing a mixture of antibiotics, constituting 1.0 g of streptomycin sulfate, 1.0 g of penicillin G potassium, 1.0 g kanamycin sulfate, 1.2 mg nalidixic acid, 125 mg vancomycin hydrochloride. After 24 h incubation, the axenic *G. lemaneiformis* was washed three times using sterile seawater to remove the antibiotics and then divided into three groups, each in triplicate. One group contained the UPB at the concentration of 2.5×10^8^ cells/ml as Group-1 (experimental group), and the other without UPB as control. The CH_4_^15N^_2_O with the concentration of 75 μmol·L^-1^ was used as the only nitrogen source in this study. The bacterial concentration required for this study was 2.5×10^8^ cells/mL determined by measured value of bacterial OD_600_ (35).

### 2.7. Measurement of urea content in medium

The urea content in culture media was measured according to the method described by Revilla et al. (36). Briefly, for urea content measurements, 0.0288 mL of color development reagent and 0.144 mL mixed reagent Ⅰ were added to 1.0 mL water sample in a 1.5 mL centrifuge tube. The color development reagent consisted of 1:1 mixed color reagent (3.0 g diacetylmonoxime and 35 mg aminourea hydrochloride were dissolved in 50 mL 50% [v/v] ethanol), and mixed reagentⅡ(20 g MnCl_2_·4H_2_O and 0.4 g KNO_3_ were dissolved in 50 mL ddH_2_O). The mixed reagentⅠconsisted of 1:20 phosphate buffer (8 g NaH_2_PO_4_·2H_2_O was dissolved in 5 mL ddH_2_O), and 100 mL H_2_SO_4_. Tubes were incubated at 70℃for 2 h. The samples were cooled for 5 min on ice, and the A_520_ was determined with a UV-visible spectrophotometer (UV2400, Sunny Hengping Instrument). The standard stock solution was prepared by dissolving 0.6 g urea in 100 mL dd H_2_O (0.1 M urea). The urea standard curve was drawn by using urea standard working solution (100 μM).

### 2.8. The content of NH_4_^+^, urea, and total cellular nitrogen in *G. lemaneiformis*

The NH_**4**_^**+**^content in *G. lemaneiformis* was spectrophotometrically analyzed as previously described by Meng et al. (37). 100 mg powdered samples were extracted in 1 mL 100 mM HCl, and subsequently, 500 μl chloroform were added to each tube. The phase was separated by centrifugation (12,000 *g*, 10 min, 8℃) after rotating for15 min at 4℃. The aqueous phase was transferred to a new 2 mL centrifuge tube containing 0.05 g activated charcoal, thoroughly mixed, and centrifuged at 20,000 *g* for 10 min at 8℃. The supernatant was transferred in a new 2 mL centrifuge tube for determining. 20 μl of the sample was mixed with 100 μl of phenol-sodium nitroprusside solution (1% [w/v] phenol and 0.005% [w/v] sodium nitroprusside were dissolved in water) and 100 μl of sodium hypochlorite-sodium hydroxide solution (1% [v/v] sodium hypochlorite and 0.5% [w/v] sodium hydroxide were dissolved in water). The mixture were incubated at 37℃for 30 min, and light absorption was measured at 620 nm. The standard stock solution was prepared by dissolving 0.5349 g NH_4_Cl in 100 mL dd H_2_O (100 μM NH_4_Cl). The NH_**4**_^**+**^standard curve was drawn by using NH_4_Cl standard working solution (1 μM).

The urea in *G. lemaneiformis* was extracted using the methods as described by Mérigout et al. (38). Briefly, 1.5 mL of 10 mM ice-cold formic acid was added to about 0.1 g of ground fresh sample. Each extract was vortexed for 15 min at 4℃and centrifuged at 16,300 *g* for 15 min at 4℃, and the supernatant was transferred in a new 1.5 mL centrifuge tube. The determination of urea content in *G. lemaneiformis* followed the above methods in measurement of urea content in media.

For determining the total cellular nitrogen in *G. lemaneiformis* we followed the methods described by Barbarino & Lourenço (39). Briefly, about 2 mg dried powdered samples of macorlage were weighed in small tin capsules and subject to combustion at 1,150℃for about 2 min in the combustion tube of a VarioEL/MICROcube elemental analyzer (Elementar Analysensysteme GmbH, Germany). The current pressure of oxygen pressure reducing valve and helium pressure reducing valve were 0.2, and 0.12 MPa, respectively. The reduction tube temperature was set at 850℃. The temperature of CO_2_ and H_2_O desorption column were set 20℃. The values were registered automatically by the recorder and integrator coupled to the analyzer.

### 2.9. Isotope analysis of δ^15^N (‰, Atm-N_2_) in *G. lemaneiformis*

The frozen mixed ball mill (Retsch MM400, Verder Group) was used to grind the algal tissues of *G. lemaneiformis*, and the powder samples after grinding were passed through a 100-mesh sieve. The pending samples and working standard samples were weighed accurately by ultra-micro analytical balance (XP6, Mettler Toledo), and were packed in the tin cup carefully. Subsequently, the tin cups containing pending samples or working standard samples were put into the automatic sampler plate in sequence. The pending samples or working standard samples were burned at 1000℃in the elemental analyzer (HT2000, ThermoFisher) to produce N_2_. The ^15^N and ^14^N ratios of N_2_ were detected by isotope ratio mass spectrometer (Delta V Advantage, ThermoFisher), and the δ^15^N values of the pending samples were calculated after comparing with the international standard (Atm-N_2_). The total nitrogen content (N%) of the pending samples was calculated by comparing the peak area of pending samples with three working standard samples.

### 2.10. Data statistics and analysis

Statistical analysis of physiological parameters among the groups was performed by repeated measures analysis of variance (5). Multivariate analysis of variance was used for statistical analysis of physiological parameters at each time point among the groups. Calculation and statistical analyses were performed with SPSS 20.0 software, with the significant threshold set to 0.05. All data were calculated with three biological replicates and reported as the means ± SD.

## 3. Results

### 3.1. Identification of bacterial isolates

A total of 41 single colonies were screened from *G. lemaneiformis, P. haitanensis*, and seawater, among which 34 bacterial isolates were identified based on 16S rDNA gene. As showed in **Table 2**, at the phylum level, these isolates were classified into Proteobacteria (76.47%), Bacteroidetes (11.76%), Firmicutes (8.82%), and Actinobacteria (2.94%).

**Table 1.**
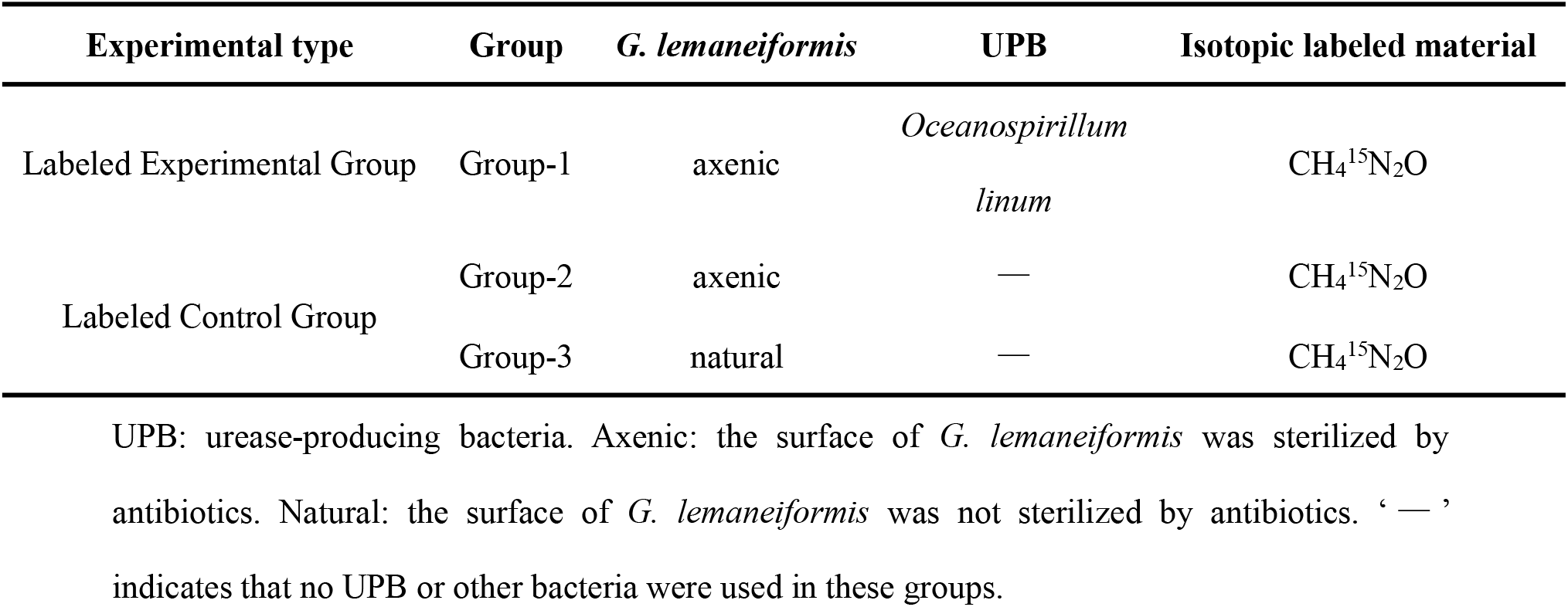
The design of co-culture experiment.

**Table 2.**
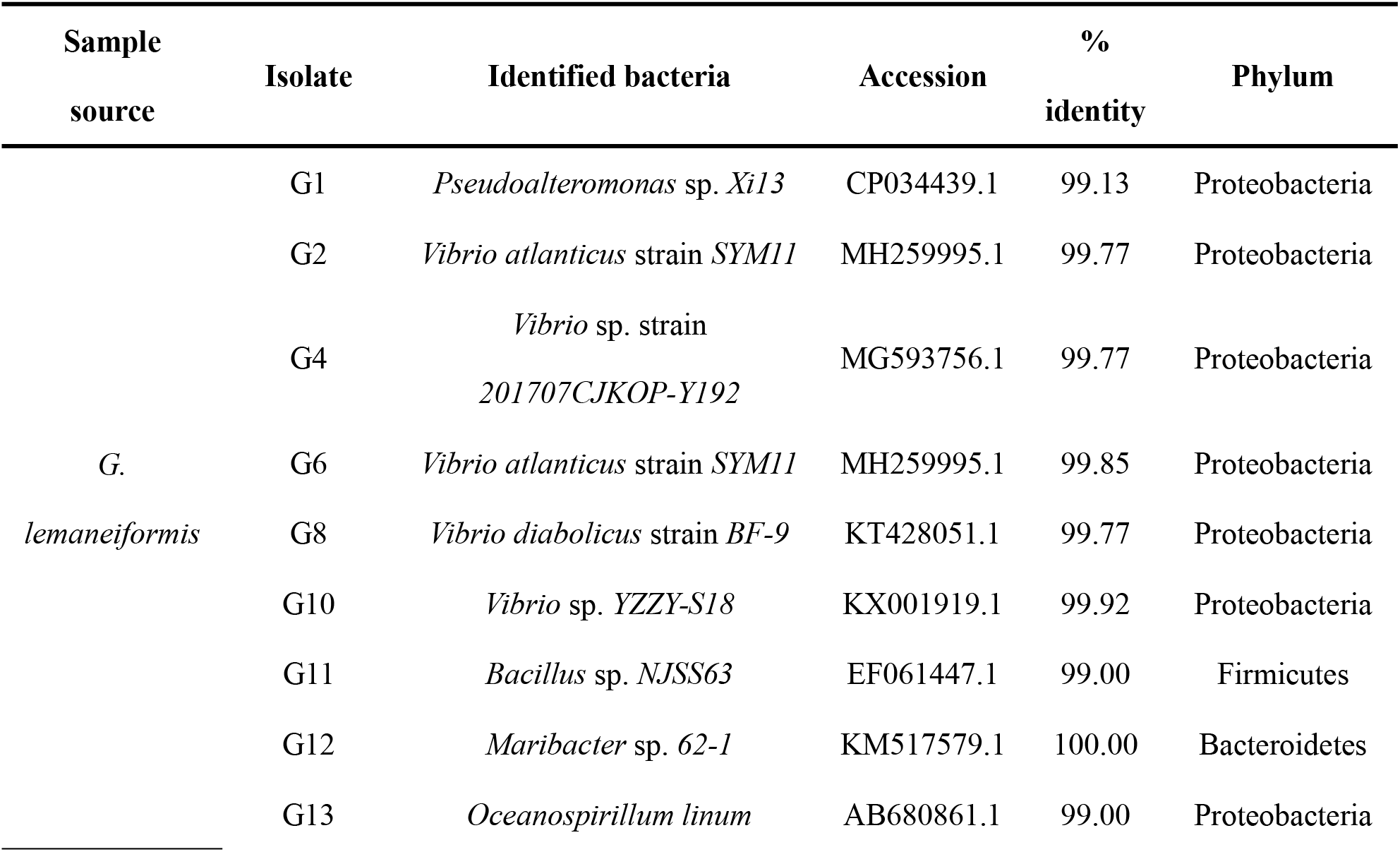
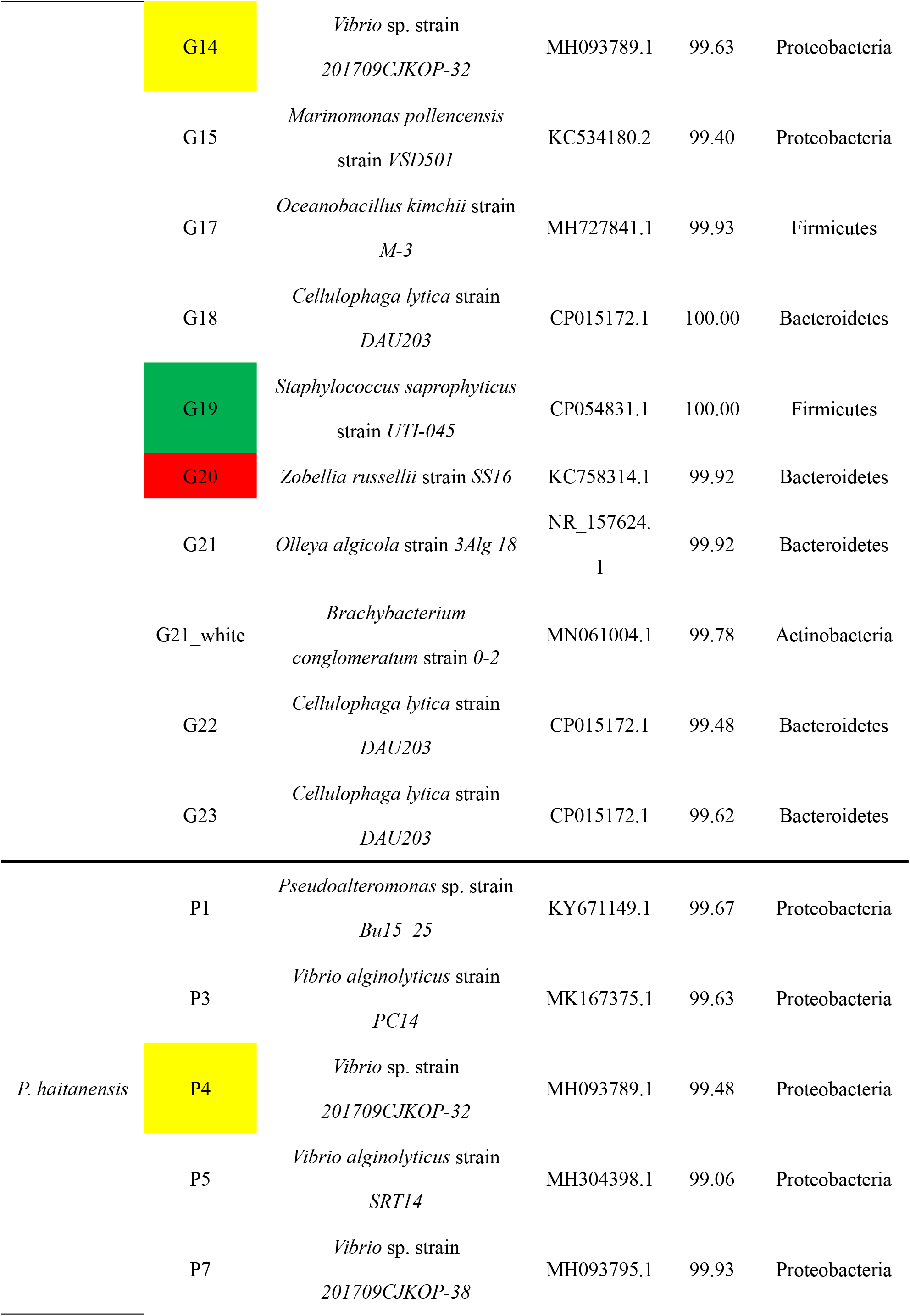
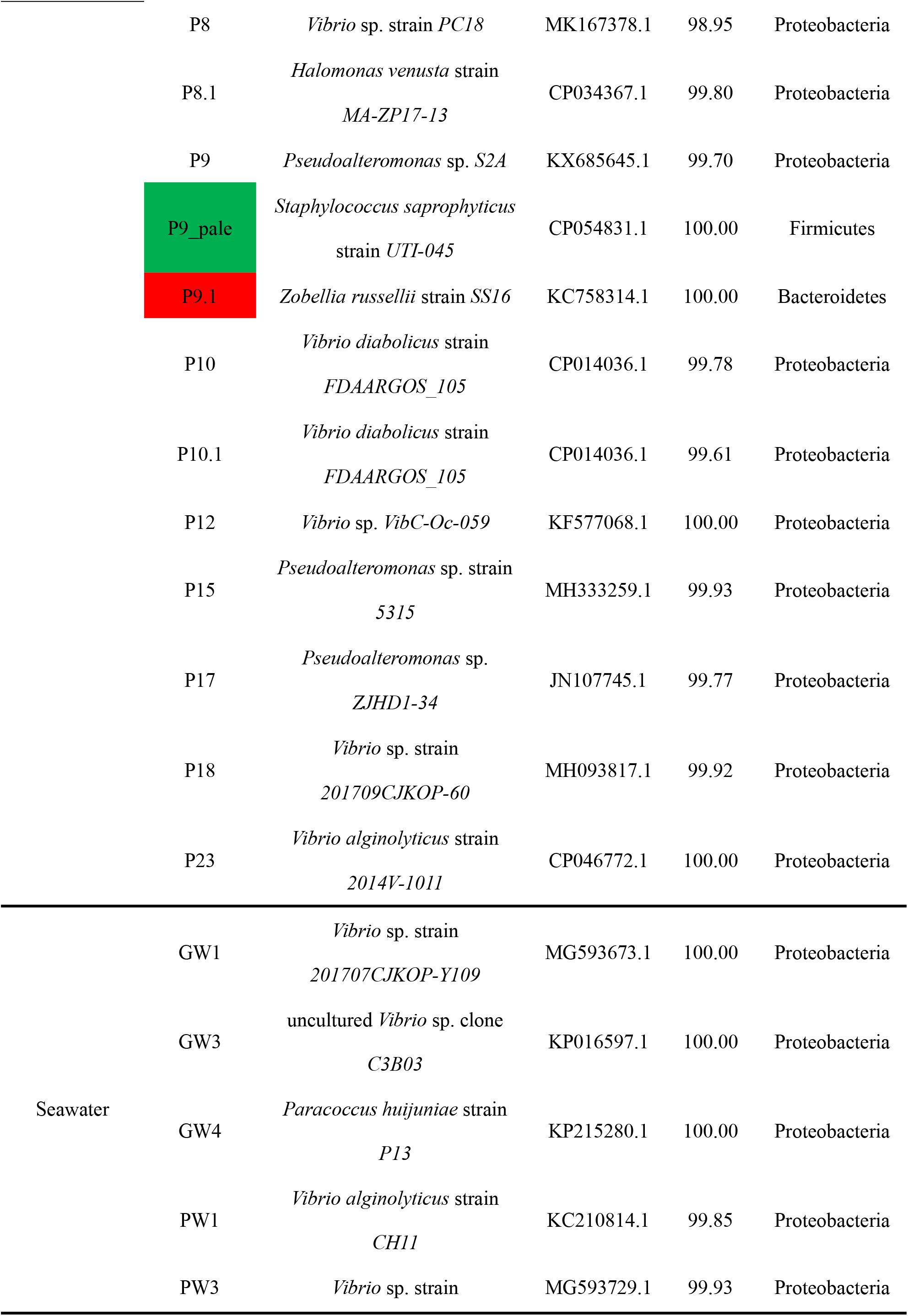
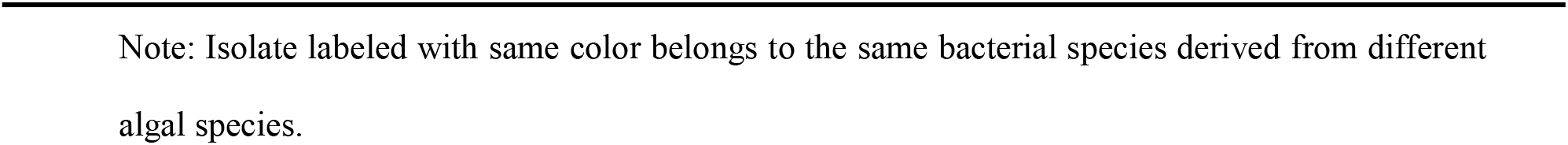
16S rRNA gene sequence identity of thirty-four bacterial isolates obtained from *G. lemaneiformis, P. haitanensis*, and seawater.

### 3.2. Screening of UPB

Totally, 12 bacteria identified as having the ability to break down urea. They are G8, G13, G15, G19, G21_white, GW3, P4, P8.1, P9, P9_pale, P12, and P23 (**Fig. 1**). Apparently, the cultivable bacteria in the cultivation environment of *G. lemaneiformis* might contain urease, which we called the potential UPB. In a previous study (20), a bacterium *Marinobacter litoralis*, which was isolated from the sponge, showed similar color change in the urea agar chromogenic plate.

**Figure 1.**
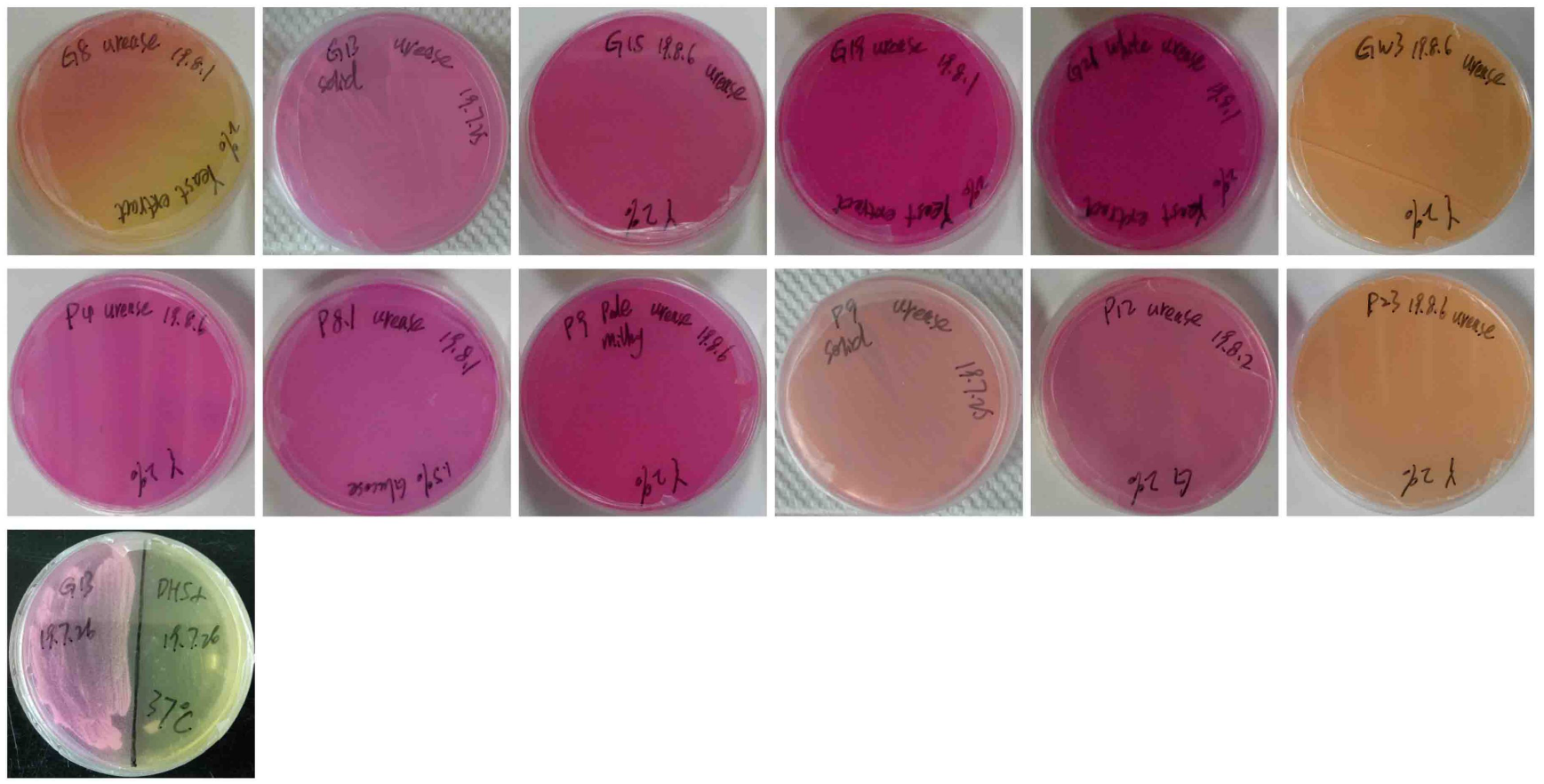
Chromogenic plate identification of UPB from the cultivation environment of *G. lemaneiformis*. DH5α: negative control.

### 3.3. *ureC* genes in UPB

*ureC* was a target gene for analysis of urea-decomposing bacteria in various environments. The *ureC* gene target band of ten potential UPB (G21_white, G19, G15, GW3, G13, P4, P9_pale, P8.1, P12, and P23) amplified by PCR was about 400 bp (**Fig. 2A & 2B**), which was consistent with the expected result. It was preliminarily believed that these strains with targeted bands contained *ureC* gene. To further verify the presence of *ureC* gene in these strains, 20 μL PCR product was taken and sequenced with L_2_F forward primer. Sequencing results showed that eight strains (G21_white, G19, G15, GW3, G13, P4, P8.1, and P12) were successfully sequenced and while two strains failed to be sequenced. Therefore, these eight potential strains were formally considered as UPB. For sequence details see Supplementary Materials (**Tab. S1**).

**Figure 2.**
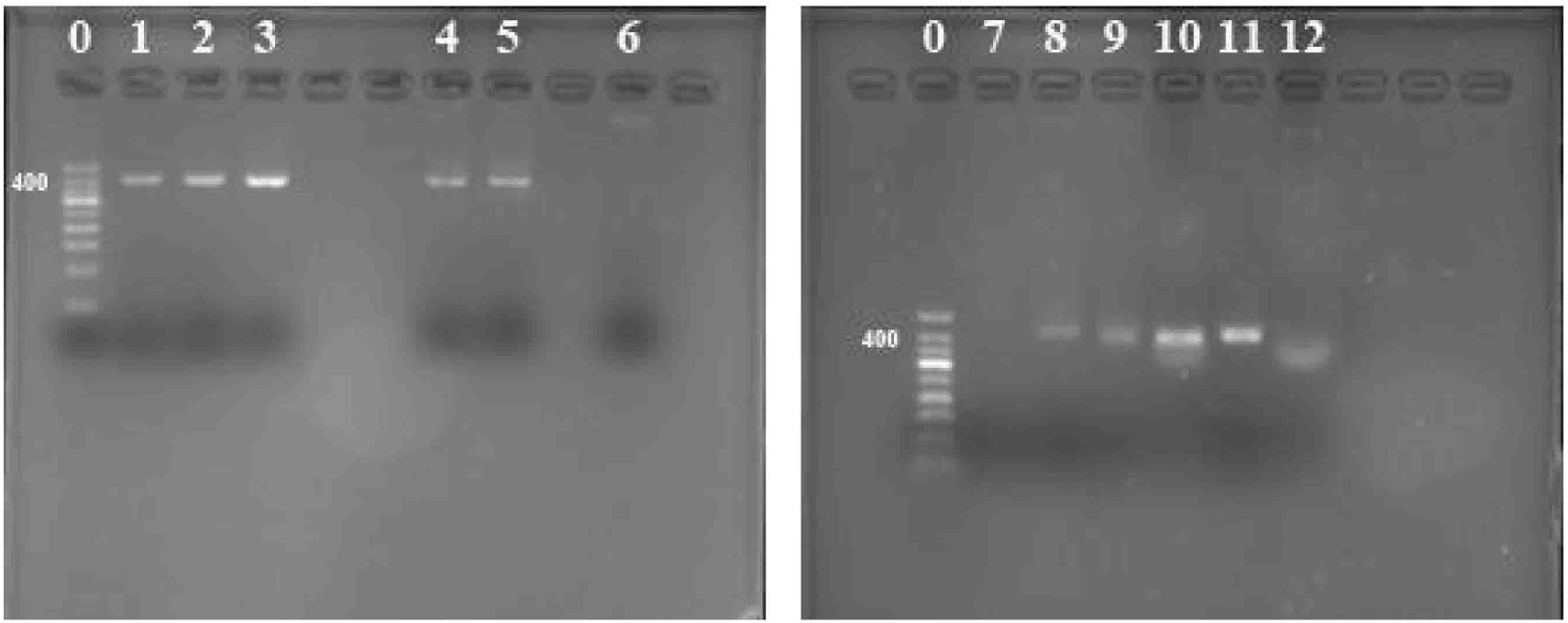
The agarose gel electrophoresis of amplified *ureC* products by direct PCR. 0: 50 bp DNA Ladder; 1-12 are strain G21_white, G19, G15, GW3, G13, P9, G8, P4, P9_pale, P8.1, P12, and P23, respectively.

### 3.4. Characteristic of urease activity from UPB

The urease activities of different UPB from *G. lemaneiformis, P. haitanensis*, and seawater are shown in **Fig. 3**. In the epiphytic bacteria associated with *G. lemaneiformis*, the urease activities of three UPB (G13, G19, and G21) were all higher than 10 U except G15 (*Marinomonas pollencensis*). However, in the epiphytic bacteria associated with *P. haitanensis*, the urease activities of all UPB (P4, P8.1, and P12) were all lower than 1 U. The strain with the highest urease activity was GW3 (34.27±4.81 U), which was isolated from surrounding seawater of *G. lemaneiformis*.

**Figure 3.**
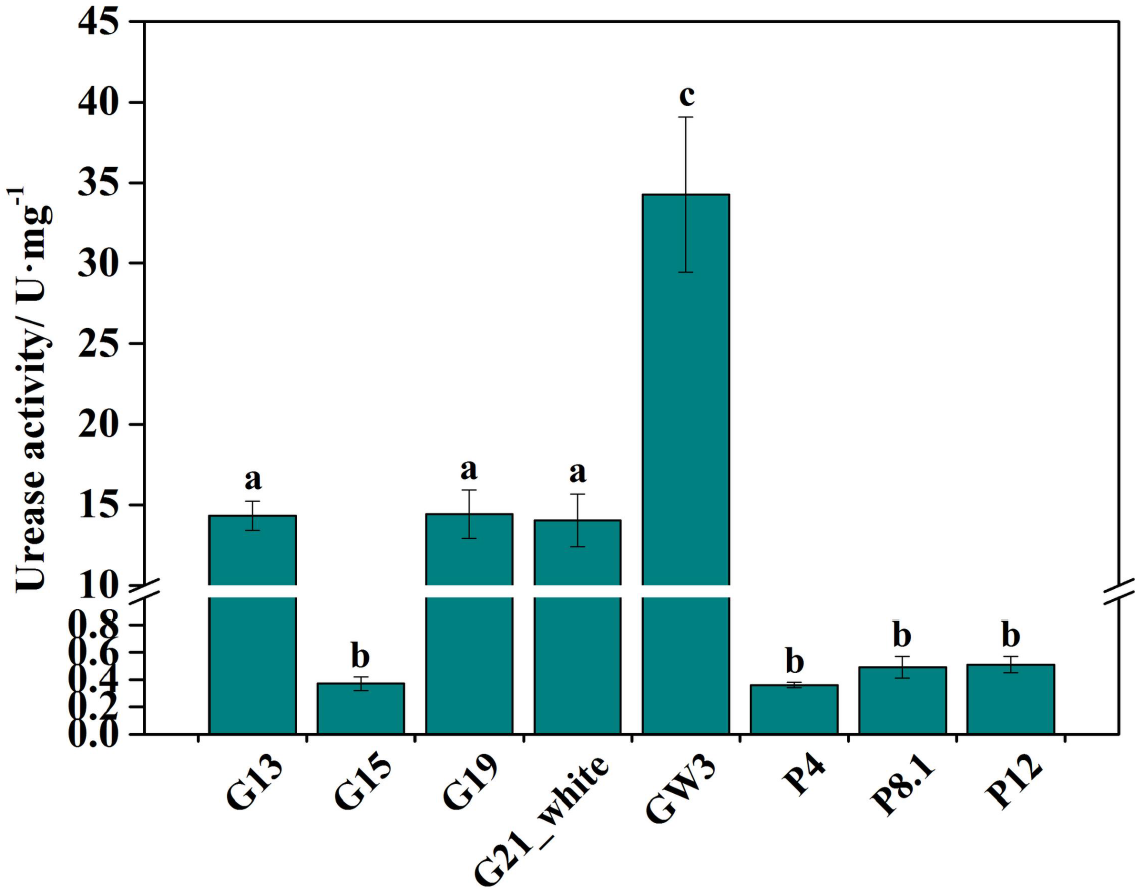
The urease activity of UPB from *G. lemaneiformis, P. haitanensis*, and seawater. Different letters (a, b, c) denote significant (*p* < 0.05) differences in urease activity between UPB.

### 3.5. Absorption rate of urea in *G. lemaneiformis*

To understand the uptake of of nitrogen from media, absorption rate of urea in *G. lemaneiformis* were measured throughout the experiment (**Fig. 4**). During whole culture period, the maximum absorption rates of urea in Group-1 (sterilized macroalage with UPB, Group-2 (sterilized macroalgae without UPB), and Group-3 (macroalgae with all epiphytes) were 5.379, 3.046, and 6.663 μmol/(g·h), respectively. There were no significant differences in urea content of culture media between Group-1 and Group-2 on 1 d and 2 d, indicating that the absorption rate of urea in *G. lemaneiformis* was basically same between Group-1 and Group-2 during first 2 days. However, the urea content of medium in Group-1 was significantly (*p* < 0.01) lower than that in Group-2 on 3 d, demonstrating that the absorption rate of urea in Group-1 was significantly (*p* < 0.01) higher than that in Group-2 at 2-3 d. Moreover, the urea content of medium in Group-3 was significantly (*p* < 0.01) lower than that in Group-1 and Group-2 during whole culture periods, showed that the absorption rate of urea in Group-3 was significantly (*p* < 0.01) higher than that in Group-1 and Group-2 at 0-3 d.

**Figure 4.**
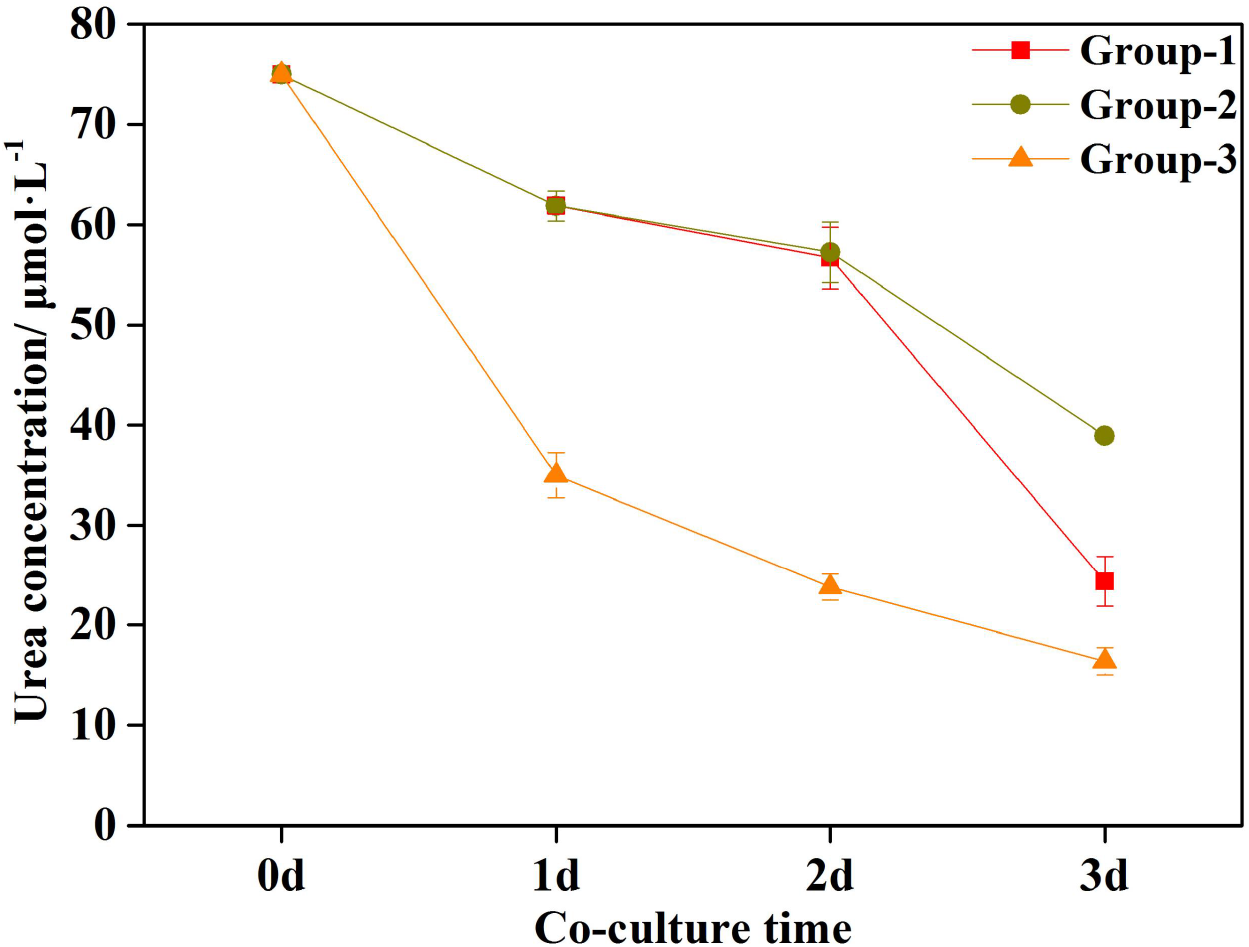
Dynamic absorption rate of urea in *G. lemaneiformis*. Group-1: axenic *G. lemaneiformis* with UPB in urea culture; Group-2: axenic *G. lemaneiformis* without UPB in urea culture; Group-3: natural *G. lemaneiformis* without UPB in urea culture. The initial urea concentration of Group-1, Group-2, and Group-3 was 75 μmol·L^-1^.

### 3.6. Physiological parameters of *G. lemaneiformis* cultured in different conditions

To investigate the response of urea absorption by *G. lemaneiformis* to UPB, some key physiological parameters were measured throughout the experiment (**Fig. 5**). The NH_**4**_^**+**^of *G. lemaneiformis* in Group-1, Group-2, and Group-3 gradually decreased with culture time. The NH_**4**_^**+**^of *G. lemaneiformis* in Group-1 was significantly higher than that in Group-2 (*p* < 0.001) and Group-3 (*p* < 0.01) on 1 d. On 2 d and 3 d, the NH_**4**_^**+**^of *G. lemaneiformis* in Group-2 was significantly (*p* < 0.01) lower than that in Group-1 and Group-3. The urea of *G. lemaneiformis* in Group-1, Group-2, and Group-3 increased to a peak firstly and then decreased. There were no significant differences in urea of *G. lemaneiformis* among Group-1, Group-2, and Group-3 on 1 d. On 2 d, the urea of *G. lemaneiformis* in Group-2 was significantly higher than that in Group-1 (*p* < 0.01) and Group-3 (*p* < 0.05). Besides, the urea of *G. lemaneiformis* in Group-3 was significantly lower than that in Group-1 (*p* < 0.05) and Group-2 (*p* < 0.01) on 3 d. The variance analysis of groups showed that the total cellular nitrogen in *G. lemaneiformis* were significantly different (*p* < 0.01) among Group-1, Group-2, and Group-3. The total cellular nitrogen of *G. lemaneiformis* in Group-1 was significantly (*p* < 0.05) higher than that in Group-2 and Group-3 during whole culture periods.

**Figure 5.**
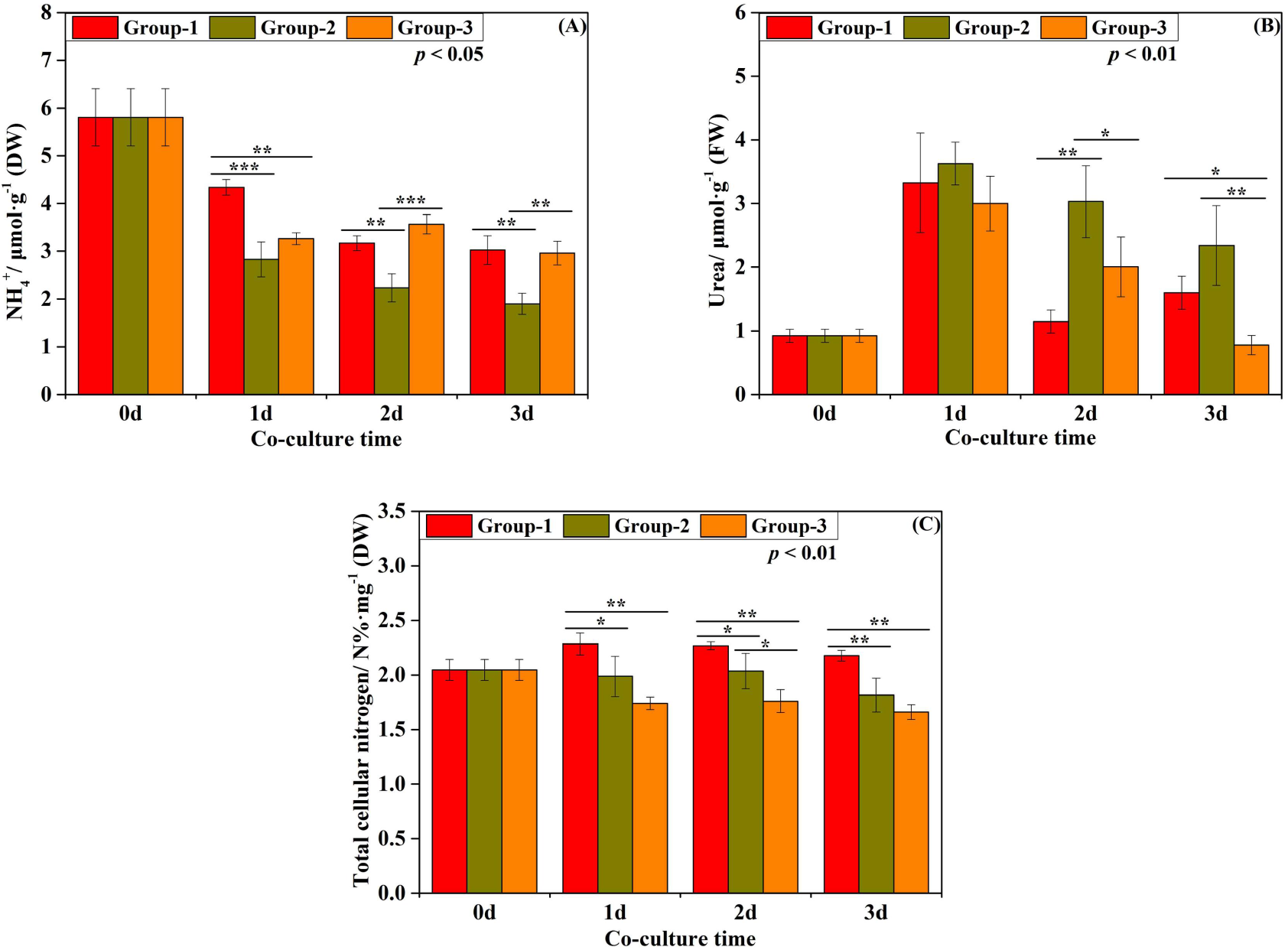
Physiological parameters of *G. lemaneiformis*. Group-1: axenic *G. lemaneiformis* with UPB in urea culture; Group-2: axenic *G. lemaneiformis* without UPB in urea culture; Group-3: natural *G. lemaneiformis* without UPB in urea culture. A: NH^4 +^ content in *G. lemaneiformis*; B: urea content in *G. lemaneiformis*; C: total cellular nitrogen content in *G. lemaneiformis*. In all experiments, error bars indicate SD in three biological replicates and asterisks indicate significance between Group-1, Group-2, and Group-3. ‘*’ represents *p* < 0.05; ‘**’ represents *p* < 0.01; ‘***’ represents *p* < 0.001.

### 3.7. Stable isotopic analysis of δ^15^N in *G. lemaneiformis*

Evidence that UPB or epiphytic bacteria associated with *G. lemaneiformis* facilitate algae uptake of δ^15^N derived from urea was provided by our isotopic analysis results (**Fig. 6**). The variance analysis of groups showed that the δ^15^N accumulation in *G. lemaneiformis* were considerably different (*p* < 0.001) among Group-1, Group-2, and Group-3. At 1 d and 2 d, the δ^15^N in *G. lemaneiformis* of Group-2 was significantly lower than that in Group-1 (*p* < 0.05) and Group-3 (*p* < 0.001). It indicated that the δ^15^N accumulation was greater in *G. lemaneiformis* with UPB (Group-1) compared to those where microorganisms had been removed (Group-2), with levels 1.20 and 1.40 times higher in *G. lemaneiformis* with UPB by 1 d and 2 d (*p* < 0.05). Similarly, the δ^15^N accumulation was greater in *G. lemaneiformis* with epiphytic bacteria (Group-3) compared to those where microorganisms had been removed (Group-2), with levels 3.99 and 3.69 times higher in *G. lemaneiformis* with epiphytic bacteria by 1 d and 2 d (*p* < 0.001). At 3 d, *G. lemaneiformis* of Group-3 had the highest (*p* < 0.01) δ^15^N value compared with Group-1 and Group-2. It demonstrated that the δ^15^N accumulation was greater in *G. lemaneiformis* with epiphytic bacteria (Group-3) compared to those where microorganisms had been removed (Group-2), with levels 1.89 times higher in *G. lemaneiformis* with epiphytic bacteria by 3 d (*p* < 0.01).

**Figure 6.**
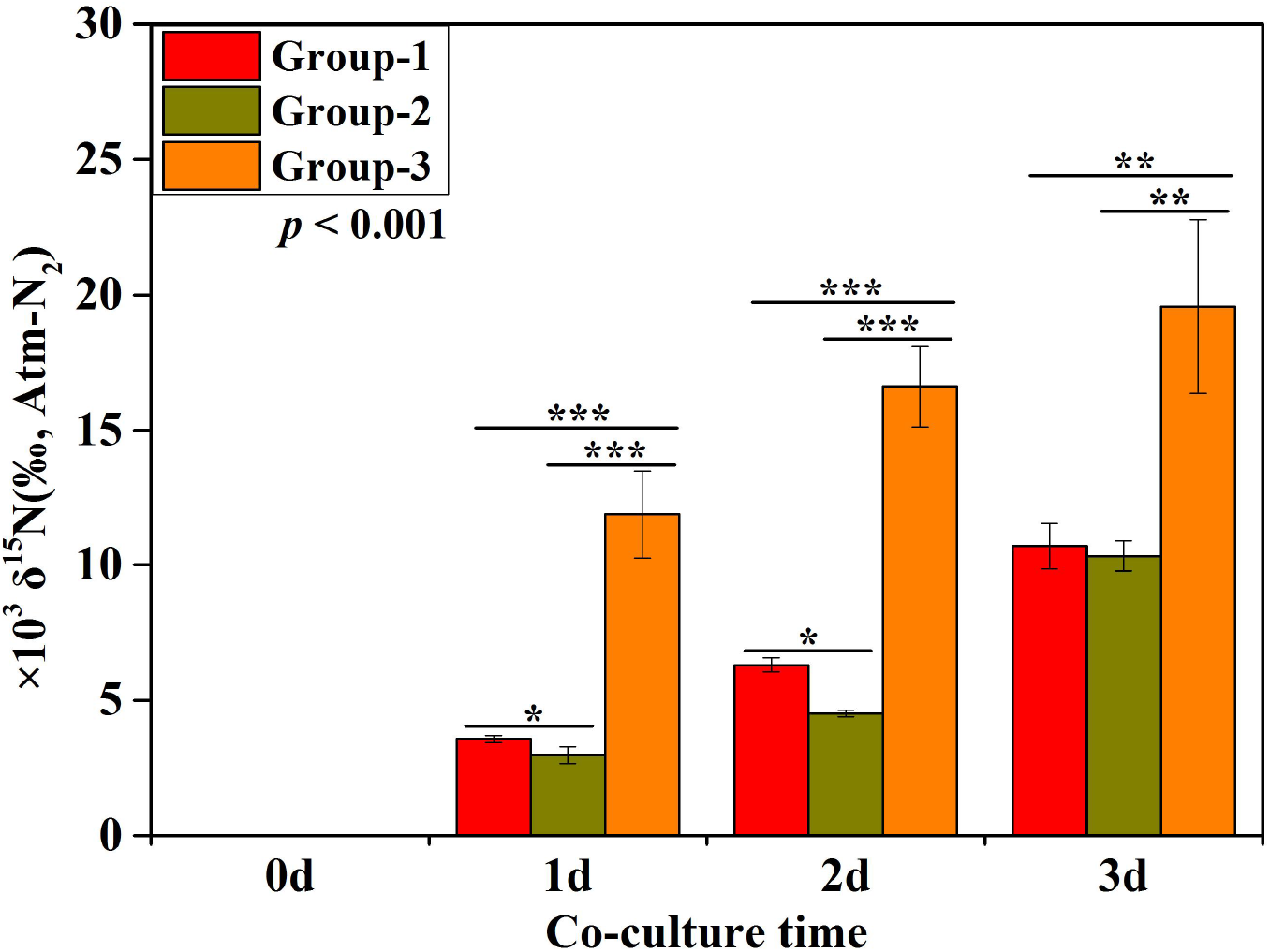
Mean (± standard deviation) δ^15^N isotopic values in *G. lemaneiformis* cultured in different conditions. Group-1: axenic *G. lemaneiformis* with UPB in urea culture; Group-2: axenic *G. lemaneiformis* without UPB in urea culture; Group-3: natural *G. lemaneiformis* without UPB in urea culture. In all experiments, error bars indicate SD in three biological replicates and asterisks indicate significance between Group-1, Group-2, and Group-3. ‘*’ represents *p* < 0.05; ‘**’ represents *p* < 0.01; ‘***’ represents *p* < 0.001.

## 4. Discussion

### 4.1. The diversity of culturable bacteria and UPB

In the current study, it was observed that higher number of bacteria were isolated from the cultivation environment of *G. lemaneiformis*. Notably, the greater number of isolates were from the surface of macroalgae rather than seawater. Similar findings have previously been reported by Ismail et al. (40), where the number of strains isolated from seaweed capable of growth on Marine Agar (MA) was significantly higher than those isolated from the surrounding seawater. Therefore, we assume that bacteria associated with algae grow better on artificial media than planktonic bacteria. This hypothesis is supported by Jensen and Fenical (41) who supposed that the adaptations of bacteria to algal surface may enhance their ability to form colonies on artificial media. Moreover, another study points out that the survival strategies evolved by seawater bacteria, including responses starvation, may dramatically reduce their ability to form colonies on nutrient-rich agar media (42). In our study, most strains isolated from macroalgae and seawater belong to Proteobacteria, which indicates that the algae-associated communities and the planktonic communities were dominated by Proteobacteria (43). The Gamma-Proteobacteria were consistently recovered from the algae-associated community and were dominated by the *Vibrio*. This suggests that *Vibrio* belongs to the resident flora of *G. lemaneiformis* and *P. haitanensis*. Interestingly, 35.29% of the strains isolated from the surface of *G. lemaneiformis* belong to Bacteroidetes. Our findings are consistent with the previous study by Pei et al. (18), where Proteobacteria (61.73%) was the most predominant phylum and Bacteroidetes (30.14%) was second dominant phylum on the surfaces of *G. lemaneiformis* in Nan’ao Island. In the present study, twelve (41.38%) of the total 29 isolates were determined to be positive for urease activity and eight isolates were formally identified as UPB by detecting *ureC* gene. To the best of our knowledge, this is the first study that used the urea agar chromogenic medium to isolate the culturable bacteria with urease activity from the cultivation environment of *G. lemaneiformis*. Moreover, the higher proportion (more than 25%) of UPB suggesting that there were a large number of UPB in the cultivation environment of *G. lemaneiformis*. UPB are widely found in various marine environment, such as in seawater (20, 22), marine sediments (44) and sponge surfaces (19). These UPB affiliate with different genera like *Bacillus, Helicobacter, Proteus, Enterobacter, Streptococcus, Escherichia, Lactobacillus*, etc (22, 23, 45). In our current study, UPB isolated from the cultivation environment of *G. lemaneiformis* belong to *Marinomonas, Staphylococcus, Brachybacterium, Halomonas*, and *Vibrio*, which indicates their environmental specificity. It can be seen that UPB are of rich and diverse species and play an indispensable role in the biogeochemical cycle by converting organic nitrogen (i.e. urea) into inorganic nitrogen (i.e. ammonium). Therefore, it can be conceived that diverse UPB in the cultivation environment of *G. lemaneiformis* play an important role in the nitrogen cycle of the coastal marine ecosystem.

The screening of potential UPB was based on the color change of urea agar chromogenic medium. However, increase in pH due to hydrolysis of peptones or other proteins and release of excessive amino acid residues in the media can also result in false positive reactions (45). It is crucial to choose a particular functional gene for urease analysis in order to further confirm whether these potential UPB indeed contain urease (46). Studies have shown that the structures of urease gene clusters from different bacterial sources differ in the number and order of their genes (47, 48). Bacterial ureases typically contain three structural genes (encoded by *ureA, ureB*, and *ureC*) and four accessory proteins (encoded by *ureD, ureE, ureF*, and *ureG*) (49).

Among them, the *ureC* gene is the largest gene encoding urease functional subunits and contains several highly conserved regions that are suitable as PCR priming sites (46). Thus, it was chosen as functional gene for urease analysis. Our data showed that eight out of twelve potential UPB contained *ureC*. The other four strains without the *ureC* gene were able to hydrolyze urea normally, suggesting that other bio-enzyme e.g. urea amidolyase (UA), which is comprised of urea carboxylase (UC) and allophanate hydrolase (AH). These are widely distributed in fungi, bacteria, and other microorganisms, and play important roles in nitrogen cycling of the biosphere (21, 50, 51). Phylogenetic analysis constructed by Strope et al. (52) showed that sequences of bacterial UA and UC were existed in *Marinomonas* and *Burkholderia*. A previous study has showed that *Marinomonas* and *Burkholderia* were detected in epiphytic bacteria on *G. lemaneiformis* by 16S amplicon sequencing (18). These results further suggested that the four strains without urease were likely to contain UA. In addition, Kanamori et al. (21) reported that there were two systems of urea degradation in bacteria, a traditional pathway catalyzed by the ureases and the UA pathway identified in their study. UC uses the energy generated by ATP decomposition to catalyze hydrolysis of urea to allophanate, which is further hydrolyzed to carbonic acid and ammonium by AH (21, 53).

The urease activities of UPB at 24 h incubation period were shown in **Fig. 3**. Our study showed that urease activity was obviously different among various strains. Bacterial urease proteins generally include structural proteins and accessory proteins (54). Structural proteins are mainly encoded by *ureA, ureB* and *ureC* genes. However, the trimer structure formed by the γ, β, and α subunits encoded by the *ureA, ureB*, and *ureC* genes was Apo-urease and didn’t have urease activity (49). Therefore, the activation of most bacterial urease requires the assistance of several accessory proteins (54), which are encoded by *ureD/H, ureF, ureG*, and *ureE* genes (49). In the present study, urease activity could hardly be detected in some UPB (G15, P4, P8.1, and P12) isolated from *G. lemaneiformis* and *P. haitanensis*, which was probably due to the lack of some accessory proteins to activate urease. The urease activities of other four UPB strains (G13, G19, G21_white, GW3) were different, which were probably related to urease structure. Studies have shown that there are some differences in the variety and quantity of accessory proteins in bacteria, which resulted in differences in structure of bacterial urease (48, 55). On the other hand, urease synthesis in some bacteria is regulated by environmental factors, such as urea and nitrogen concentrations or pH (56, 57). Nitrogen availability can influence the urease activity of bacteria by modulating *ureC* gene transcription (56). Thus, changes of various environmental factors may hinder the synthesis of bacterial urease or inhibit the expression of urease gene, and then affect the urease activity. In a word, the bacterial urease activity is not only depends on its structure but also on the environment.

### 4.2. The ecological function of UPB

To study the ecological functions of UPB, the UPB and axenic *G. lemaneiformis* were co-cultured to explore the response of urea uptake by *G. lemaneiformis* in the presence of UPB. The effect of UPB on urea uptake in *G. lemaneiformis* was evaluated by determining urea content in culture media and some other physiological parameters of macroalgae. Our data revealed that the urea content in culture media of Group-1 was significantly (*p* < 0.01) lower than that of Group-2 (**Fig. 4**), indicating that the urea consumption of Group-1 was significantly (*p* < 0.01) higher than that of Group-2. Moreover, the urea consumption in Group-3 was dramatically (*p* < 0.01) greater than that in both Group-1 and Group-2 (**Fig. 4**). Urease produced by diverse bacteria can catalyze the hydrolysis of urea to ammonium (58), which becomes the preferred nitrogen source of many organisms (59). A part of urea was decomposed by urease secreted by UPB after adding *Oceanospirillum linum* bacterium in Group-1, thus the urea consumption of Group-1 was obviously higher than that of Group-2. Most organisms and even microbial symbionts that use urea as a nitrogen source rely on urease (19). Ammonium was not only utilized by UPB, but also by *G. lemaneiformis*, which resulted in the NH_**4**_^**+**^-N content and the percentages of total N of *G. lemaneiformis* in Group-1 was dramatically (*p* < 0.05) higher than that in Group-2 (**Fig. 5A**). It’s worth noting that *G. lemaneiformis* still normally used urea without the participation of microorganisms (**Fig. 4**). Our findings are consistent with the study by Tarutani et al. (60), which direct utilization of DON by *Ulva pertusa* without associated microorganisms. Moreover, Engeland et al. (9) also proposed that even after removal of epiphytes, phytoplankton, and partial bacterial communities, macrophytes were able to absorb nitrogen immediately from organic origin. These studies provide a scientific basis for differentiating between direct DON uptake and uptake after remineralization by the bacterial community. In addition, DON can be used by many phytoplankton directly through processing mechanisms, such as urease activity and amino acid oxidation (61, 62). Our previous study (data not published) showed that UA sequences were found in the transcriptome data of *G. lemaneiformis*, indicating *G. lemaneiformis* had the potential ability to utilize organic nitrogen. Besides, the importance of organic nitrogen to macroalgal N nutrition depends on the availability of both dissolved inorganic and organic nitrogen compounds (63). In current study, *G. lemaneiformis* was cultured in organic nitrogen source (urea) after 4 days of low inorganic nitrogen culture, and urea would be the only nitrogen source left for *G. lemaneiformis* to absorb and utilize. Tyler et al. (64) have demonstrated that there were a relatively small fraction of amino acids and urea assimilated by macroalgae where the inorganic nitrogen supply in environment was high. Yet, when dissolved inorganic nitrogen was low, and organic compounds played a much more important role (64).

In addition, the ecological function of urease has not been fully evaluated at the physiological level, so it is necessary to use stable isotopic tracer technique to track the ^15^N accumulation in macroalgae. To the best of our knowledge, this is the first study that used stable isotopic tracer technique to evaluate the effects of UPB on urea utilization in *G. lemaneiformis*. Evidence that UPB associated with *G. lemaneiformis* facilitate algae uptake of ^15^N derived from urea was provided by our results (**Fig. 6**). At all times following culture, ^15^N accumulation was greater in *G. lemaneiformis* with UPB (Group-1) compared to those where microorganisms have been removed (Group-2), with levels 1.40 times higher in *G. lemaneiformis* with UPB by 2 d (*p* < 0.01). Similarly, ^15^N accumulation was greater in *G. lemaneiformis* with epiphytic microorganisms (Group-3) compared to Group-2, with levels 3.98 times higher in *G. lemaneiformis* with epiphytic microorganisms by 1 d (*p* < 0.01). These results clearly indicated that more organic nitrogen sources could be absorbed by *G. lemaneiformis* in the presence of UPB or epiphytic microorganisms. Uptake of organic nitrogen (urea or DFAA) by macroalgae is based on its competitive advantage, especially under inorganic nitrogen stress (60). Tarquinio et al. (14) reported that ^15^N accumulation was greater in leaves of seagrass with an associated microbiota compared to those where microorganisms had been removed, with levels 4.5 times higher in leaves with microorganisms by 12 h. In fact, mineralization of organic matters by seagrass associated microorganisms may increase the availability of N for uptake by seagrass (13, 15). There is evidence that using bulk isotope does not support for specific isotope tracer accumulation point (14). Therefore, high-resolution secondary ion mass spectrometry (NanoSIMS) will be required to trace the uptake of ^15^N derived from urea in future studies. The distinction of direct or indirect uptake of organic nitrogen in many phytoplankton and microbes has been well documented (8, 65) but not for macroalgae. Smith et al. (12) reported that rates of urea uptake calculated by ^15^N enrichment were approximately two fold higher than those based on ^13^C enrichment, thus demonstrating that direct uptake of the urea molecule in giant kelp. Therefore, the double isotope labeling method is of great significance for exploring the contribution ratio of microorganisms to the uptake of organic nitrogen by algal host. The direct uptake of urea molecule by macroalgae is likely benefit by a urea-transporting protein called DUR3-like (66, 67). DUR3 proteins mediate high-affinity transport of exogenous and endogenous urea (67). Based on the above reports, we assume that the macroalgae *G. lemaneiformis* can directly absorb urea by DUR3-like transporter. Decomposition of organic nitrogen by associated microorganisms, especially UPB, may enhance the availability of nitrogen for uptake by *G. lemaneiformis*.

## 5. Conclusions

Eight UPB with *ureC* gene were obtained from the cultivation environment of *G. lemaneiformis* by isolation, screening, and PCR identification. There were some differences in urease activity among these UPB, which indicated that urease produced by various UPB had different ability to degrade urea. In the algae-bacteria co-culture, physiological analysis showed that urea consumption in medium increased significantly when UPB or epiphytic bacteria were present compared with axenic culture. It was speculated that urea could be utilized by *G. lemaneiformis* without the participation of microorganisms, but urea uptake of *G. lemaneiformis* would be promoted by UPB or epiphytic bacteria. In addition, the results showed that total nitrogen content in the tissues of *G. lemaneiformis* increased significantly (*p* < 0.05) in the presence of UPB compared with axenic culture. Stable isotope labeling was used to track the accumulation of nitrogen in algae. It was found that the δ^15^N isotope values in algae increased markedly (*p* < 0.01) with the addition of UPB or epiphytic bacteria, which further revealed that urea is utilized by *G. lemaneiformis*, and UPB or epiphytic bacteria promoted urea utilization in *G. lemaneiformis*. Hence, it has been proved that *G. lemaneiformis* uses organic nitrogen, in the presence of functional microorganisms on its surface who significantly contribute to boosting its absorption.

## Data availability

The datasets presented in this study can be found online repositories, the names of the repository/repositories and accession number(s) can be found in the article/Supplementary Material.

## Acknowledgments

We thank Shenzhen Huake Jingxin Testing Technology Co., Ltd. for providing isotope testing services, and Dr. Zepan Chen of Shantou University for his help with sampling.

## Author contributions

**Pengbing Pei:** Conceptualization, Methodology, Data curation, Investigation, Writing-original draft, Writing-review & editing, Formal analysis, Visualization. **Hong Du:** Conceptualization, Resources, Writing-review & editing, Supervision, Funding acquisition, Validation, Visualization. **Muhammad Aslam:** Conceptualization, Methodology, Investigation, Writing-review & editing. **Hui Wang:** Writing-review & editing. **Peilin Ye:** Methodology, Investigation. **Tangcheng Li:** Writing-review & editing, Visualization. **Honghao Liang:** Methodology, Investigation. **Zezhi Zhang:** Methodology, Investigation. **Xiao Ke:** Methodology, Investigation. **Qi Lin:** Writing-review & editing. **Weizhou Chen:** Writing-review & editing.

## Funding

This research was supported by the National Natural Science Foundation of China (grant number 41976125), the Open Program of Key Laboratory of Cultivation and High-value Utilization of Marine Organisms in Fujian Province (2022fjscq02), the Program for University Innovation Team of Guangdong Province (2022KCXTD008), the Natural Science Foundation of China grants (grant number 42206116), and the Team Project of Department of Education of Guangdong Province (2018KCXTD012).

## Competing interests

The authors declare no competing interests.

